# Bayesian inference of neuronal ensembles

**DOI:** 10.1101/452557

**Authors:** Giovanni Diana, Thomas T. J. Sainsbury, Martin P. Meyer

**Affiliations:** Center for Developmental Neurobiology & MRC Center for Neurodevelopmental Disorders, King’s College London, Guy’s Hospital Campus, London SE1 1UL, UK

## Abstract

In many areas of the brain, both spontaneous and stimulus-evoked activity can manifest as synchronous activation of neuronal ensembles. The characterization of ensemble structure and dynamics provides important insights into how brain computations are distributed across neural networks. The proliferation of experimental techniques for recording the activity of neuronal ensembles calls for a comprehensive statistical method to describe, analyze and characterize these high dimensional datasets. Here we introduce a generative model of synchronous activity to describe spontaneously active neural ensembles. Unlike existing methods, our analysis provides a simultaneous estimation of ensemble composition, dynamics and statistical features of these neural populations, including ensemble noise and activity rate. We also introduce ensemble “coherence” as a measure of within-ensemble synchrony. We have used our method to characterize population activity throughout the tectum of larval zebrafish, allowing us to make statistical inference on the spatiotemporal organization of tectal ensembles, their composition and the logic of their interactions. We have also applied our method to functional imaging and neuropixels recordings from the mouse, allowing us to relate the activity of identified ensembles to specific behaviours such as running or changes in pupil diameter.

## Introduction

Recent advances in multineuronal recording techniques enable recording the activity of thousands of neurons simultaneously and with single neuron resolution. For instance, calcium imaging and neuropixels probes are powerful approaches to describe the spatiotemporal structure of population activity across multiple brain regions[8, 7, 25, 19, 18]. Such techniques have revealed that in many areas of the brain a prominent feature of both evoked and spontaneous activity is the organization of neurons into ensembles; groups of neurons that tend to fire in synchrony[3, 4, 15, 1, 30]. Importantly, spontaneously active ensembles are similar to those evoked by sensory stimuli suggesting that ensembles encode features of the sensory environment and that their spontaneous activation reflects an intrinsic capacity of the brain to generate an internal model of the environment[2, 30, 5]. In the zebrafish tectum ensemble activation predicts tail movements suggesting that motor variables may also be encoded by ensemble activity[30]. As a result of such studies ensembles are increasingly believed to be the units of brain computation[29, 35, 16, 30]. Characterization of ensemble structure and dynamics may therefore provide crucial insights into how brain computations are distributed across neural networks.

The problem of identifying neuronal ensembles from large scale recording is computationally challenging due to a degree of stochasticity that characterizes neuronal activity. For instance, neurons may fire independently of the ensemble of which they are a member and not all neurons within an ensemble might be recruited when the ensemble fires. Moreover, ensembles can exhibit temporal overlap, which makes it difficult to define them, especially with a limited number of recorded synchronous events. One approach to identifying ensembles is to apply dimensionality reduction techniques (PCA/SVD) followed by rotation methods (factor analysis, ICA) where neuronal populations are identified as linear “features”[30, 5, 6, 22, 11, 21]. However, these techniques require an a priori estimate of the number of ensembles which is challenging to validate with a level of statistical certainty. An alternative approach consists of representing the observed activation patterns as nodes of a graph[1]. This method provides a characterization of the neural activity in terms of repeated patterns of neurons (‘assemblies’) obtained by partitioning the graph using spectral clustering. The main drawback of these methods is that statistical confidence on ensemble properties such as size, identity of constituent neurons and dynamics cannot be determined due to the absence of an underlying model. Moreover, the uncertainty on neuronal identity is not propagated when estimating statistical features of neuronal ensembles.

The solution to this problem is to introduce Bayesian models of neuronal activity where all assumptions underlying the definition of neuronal ensembles are clearly stated. Earlier works on Bayesian estimation of neural connectivity utilized realistic models of neural dynamics such as the leaky integrate-and-fire model[24, 14]. However, the complexity of these models introduces scalability issues when applied to large networks. The inherent limitations of existing techniques mean that some basic but fundamental features of ensembles remain poorly characterized. For instance, it is not even clear what fraction of neurons in the brain can be reliably assigned to an ensemble. How many neurons does a given ensemble contain and to what extent is the activity of a neuron locked to the ensemble of which it is a member? If such features could be reliably quantified it would enable statistical estimates of how ensemble architecture and dynamics change as a function of development, brain state and behaviour, for example.

Here we introduce a different approach to analyze neuronal ensembles where Bayesian inference is used to match the observed synchrony among neurons to a generative model. Our method allows us to combine the information from all synchronous events, identifying the ensemble structure without the need for dimensionality reduction or correlation measures. In our generative model the state of the neurons depends on the state of the ensemble they belong to, introduced as latent variables in the model. Unlike existing methodologies, our method generates estimates of ensemble ON/OFF states over time, providing a coarse-grained representation of network activity. Together with ensemble dynamics our inference method can be used to determine within-ensemble statistical features such as noise, which quantifies the probability of the member neurons to fire when the ensemble is inactive, activity rate, which is the probability of the ensemble to be active, and coherence which is a measure of synchrony between constituent neurons. These biologically relevant features of ensemble dynamics have until now been challenging to quantify using existing methods.

Using our method we have characterized spontaneously active neuronal ensembles from large-scale functional imaging data obtained from zebrafish tectum and mouse visual cortex. We also demonstrate that our method can be applied to diverse data types by analyzing data from a neuropixels probe located in mouse visual cortex, hippocampus and thalamus. We demonstrate that the spatiotemporal properties of spontaneous ensembles in zebrafish tectum are remarkably stereotyped across larvae. Furthermore, we have exploited the ability to describe ensemble ON/OFF states to generate graphs that reveal functional interactions between ensembles. These demonstrate that ensembles are organised into topographically restricted subnetworks. By analyzing the functional imaging and neuropixels data from mouse we could link the activity of multiple identified ensembles to specific behavioral outputs such as running speed or changes in pupil diameter. Our method thus provides a statistically rigorous approach to identifying ensembles, their intrinsic dynamics, interactions and coordinated activation during behaviour.

## Theory

### The generative model

In this section we outline the set of generative rules to sample the activity of *N* neurons organized in *A* ensembles for *M* time frames. The ensemble membership is specified by assigning a label *t*_*i*_ to each neuron *i* as an integer between 1 to *A* drawn from a categorical distribution with probabilities *n*_*µ*_, *µ* = 1, *…, A*. In the following we will adopt the shorthand notation ***µ*** = {*i ∈ {*1, *· · ·, N*}: *t*_*i*_ = *µ*} to indicate the set of neurons assigned to the same ensemble *µ*. Vectors of parameters will be denoted by dropping the indexes, i.e. *p = {p*_1_, *· · ·, p*_*A*_}. The activity states *ω*_*kµ*_ = {0, 1} of ensemble *µ* at time *k* are independent Bernoulli latent variables with ensemble-specific probabilities *p*_*µ*_ *≡ P (ω*_*kµ*_ = 1). The activity of all neurons at all times, denoted by the binary matrix *s*_*ik*_ is drawn from the conditional probability

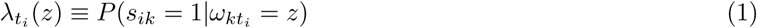

depending on the state of the corresponding ensemble *t*_*i*_. The parameters *λ*_*µ*_(1) and *λ*_*µ*_(0) represent the probabilities of any of the neurons belonging to the ensemble *µ* to fire when the ensemble is respectively active or inactive. From here onwards we will refer to *λ*_*µ*_(0) as the activity noise in the ensemble and to *λ*_*µ*_(1) as the “coherence”; the propensity of an ensemble’s constituent neurons to fire synchronously. The model parameters *θ* ={*n, p,λ*} provide a full characterization of the ensembles based on the statistical properties, including firing frequency, size, noise and coherence levels.

### Likelihood

The joint probability of neuronal membership *t*, the ensemble activity matrix *ω* and the neuronal activity matrix *s* conditional to the model parameters *θ* (likelihood) can be written as

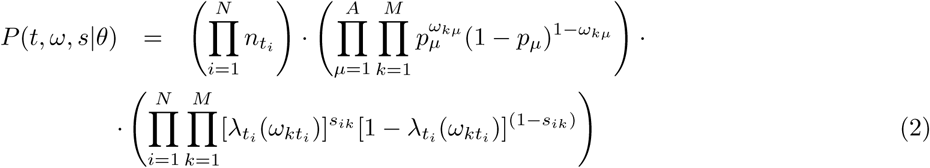

To perform Bayesian inference we need to introduce prior distributions on the model parameters. A natural choice is given by

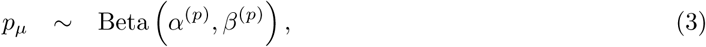

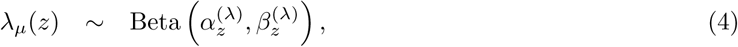

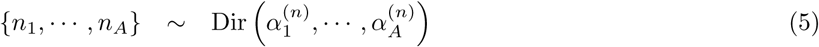

where the ensemble activity rates *p*’s and the conditional activity rates *λ*’s are drawn from a beta distribution while a Dirichlet prior is assigned to the mixture probabilities *n*’s. Where *α*’s and *β*’s are the (hyper-)parameters characterizing our prior knowledge on the model parameters. This model can be represented graphically as in Fig. 1A.

**Figure 1:**
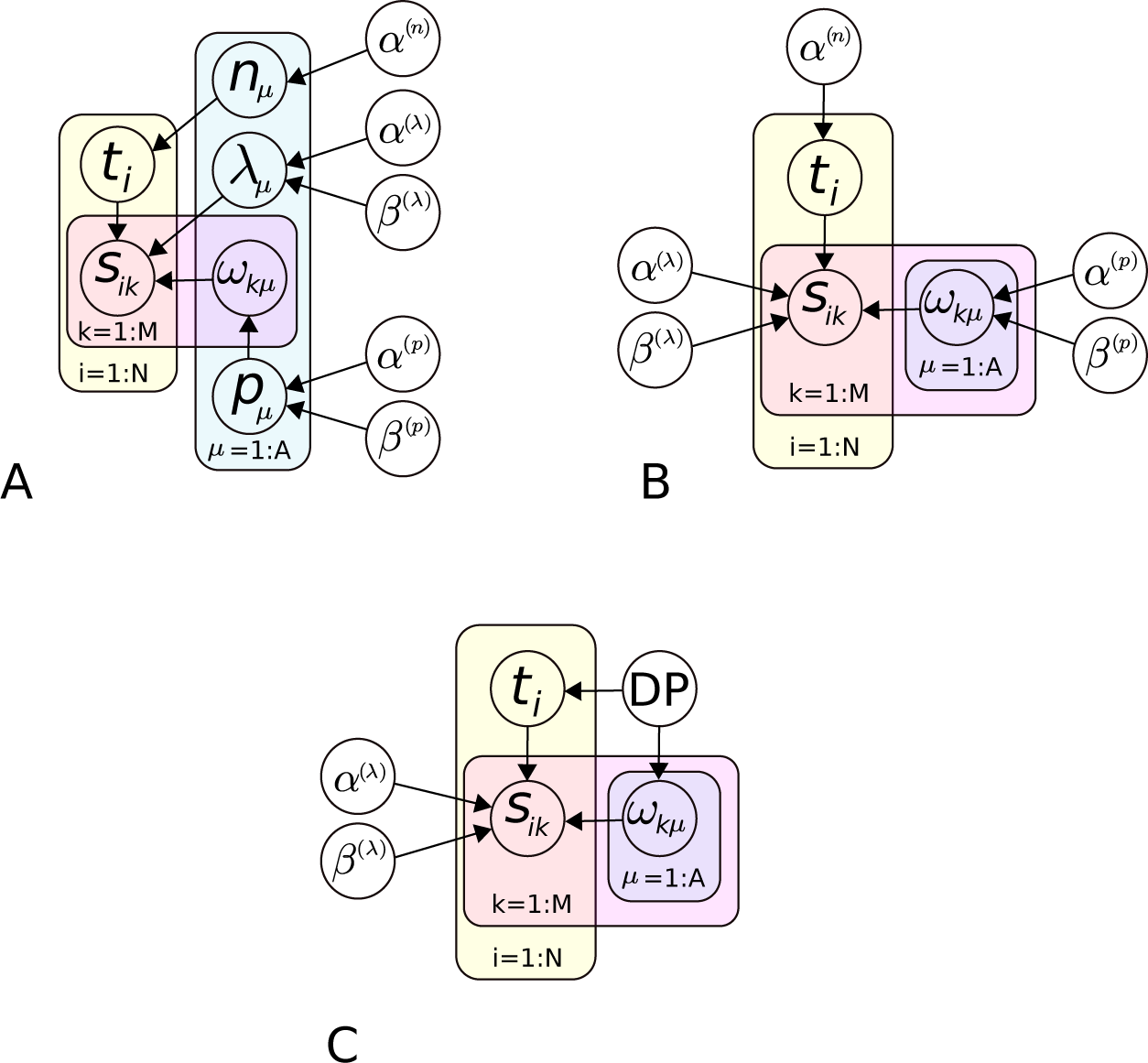
Graphical representation of the generative model. (A) Full model, (B) collapsed model with integrated parameters, (C) collapsed model with Dirichlet process prior.

The present formulation of the generative model can be used to derive a Gibbs sampler of the posterior distribution, however it would not be very efficient. To improve inference accuracy we can integrate out the parameters *θ* and obtain the marginal probability *P (t, ω, s*). This procedure leads to the “collapsed” model depicted in Fig. 1B which in general reduces the uncertainty associated to the estimation of the remaining variables. To proceed our derivation we first introduce the summary variables

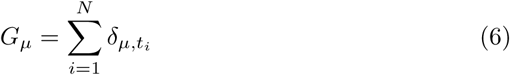

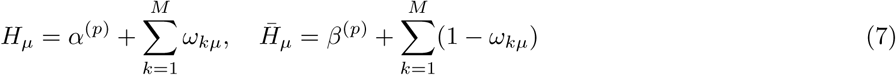

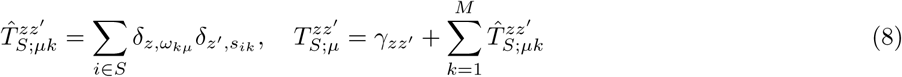

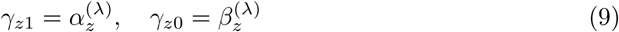

where *δ*_*ij*_ is the Kronecker delta function and *S* is any subset of indexes in the range 1 to *N*. To simplify the notation we also introduce additional definitions

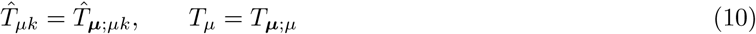

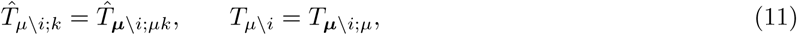

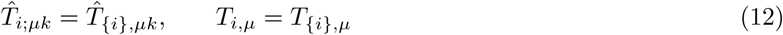

For each ensemble *µ, G*_*µ*_ is the ensemble size, *H*_*µ*_ and 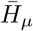 denote (up to an additive constant) the number of active and inactive events over time respectively, while the matrix 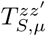counts how many times the state (*ω*_*kµ*_, *s*_*ik*_) = (*z,z’*) is observed across time and the neurons in the set *S*.

Due to the conjugate character of the prior distributions the integration over *θ* can be carried out analytically, leading to the marginal likelihood

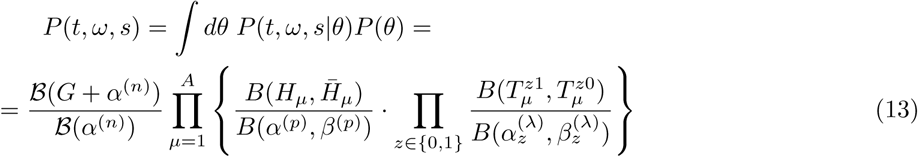

where *B*(*·, ·*) is the Euler beta function and *B* is defined as the product of gamma functions

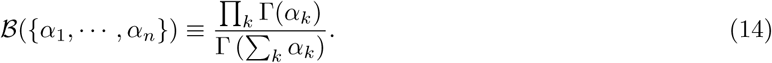

After integrating out *θ* we can rewrite the joint probability in Eq. (13) as the product between the likelihood

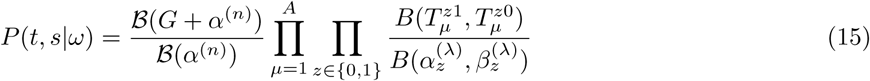

and the prior on the latent variables

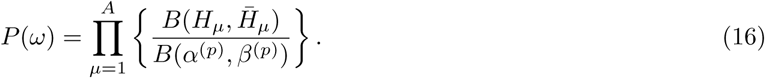

We will employ this “collapsed” formulation of the model to make inference on cell membership and ensemble activity.

### Inference

Given the data on neural activities in the form of a binary matrix *s*, we can use the generative model described above to make inference on neuronal identities and ensemble activity along with the model parameters. Within our probabilistic framework, we can estimate these quantities by evaluating their expectations with respect to the conditional distribution *P (t, ω*|*s*). These averages can be computed via Montecarlo methods by generating random samples from the joint distribution.

We first consider the case where the number of neuronal ensembles *A* is known. To obtain samples from *P (t, ω, s*) in Eq. (13) we can use a Gibbs sampler where cell membership and ensemble activities are drawn sequentially from the conditional distributions *P (t*|*ω, s*) and *P (ω*|*t, s*) respectively.

#### Algorithm 1 Collapsed Gibbs sampling

~~~
1: Initialize *t*
2: **while** convergence criteria **do**
3: **for each** ensemble *µ ∈* [1, *A*] time *k ∈* [1, *M*] **do**
4: draw *ω*_*kµ*_ *∼ P (ω*_*kµ*_|*t, s*)
5: **for each** cell *i ∈* [1, *N*] **do**
6: draw *t*_*i*_ *∼ P (t*|*ω, s*)
7: draw *θ ∼ P (θ*|*t, ω, s*)
~~~

The conditional distribution of the membership of the *i*th data point *t*_*i*_ can be written as

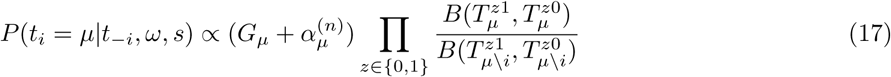

where we used the notation *t*_*-j*_ = {*t*_1_, *· · ·, t*_*j-*1_, *t*_*j*+1_, *· · ·, t*_*N*_}.

The conditional probability for the ensemble activation matrix *ω* is given by

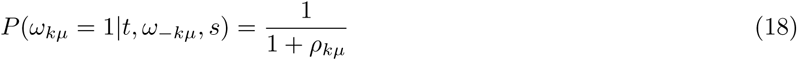

Sampling the parameters *θ* from the posterior distribution *P (θ*|*t, ω, s*) is easy thanks to our choice of the corresponding priors. The posterior distributions of *p*_*µ*_, *λ*_*µ*_ and *n*_*µ*_ retain the same functional form as in Eqs. (3,4,5) with updated hyper-parameters

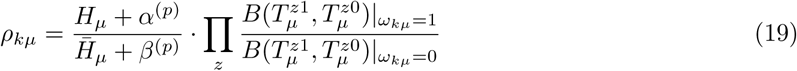

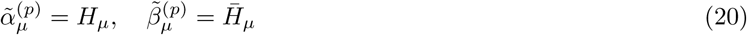

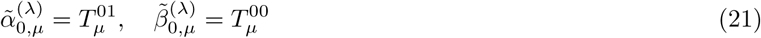

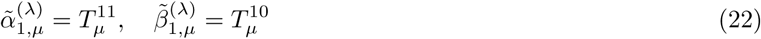

Where we have added an extra label *µ* to the posterior hyperparameters corresponding to different ensembles.

### Inference of the number of neuronal ensembles

So far we have discussed the case where the number of ensembles *A* is known. In general this information is not available, therefore we need to extend the analysis in order to make predictions on the number of neuronal ensemble present in the data. Estimating the number of components in a mixture is a common issue in unsupervised machine learning. In the context of our model we employ the Dirichlet process[13] (DP) as a prior on the mixing measure describing the composition of the data (see Fig. 1C for a graphical representation). The DP prior can be implemented by introducing specific Metropolis-Hastings acceptance rules to increase or decrease the number of components[26], which generalize the inference problem to an arbitrary number of ensembles.

For each neuron *i*, its membership *µ*^*∗*^ is drawn from the proposal

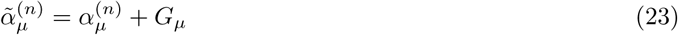

where *α* is the concentration parameter of the Dirichlet process. Existing ensembles are drawn proportionally to their occupancy 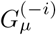 (calculated from all membership except the current *µ*_*0*_). If a new ensemble is proposed, *µ* = *A*+1, then we draw a corresponding binary vector *ω*_*kµ*_ from the prior distribution in Eq. (16). The proposal is accepted with probability

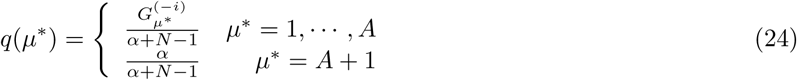

#### Algorithm 2 Metropolis-Hastings sampling with Dirichlet process prior

~~~
Initialize *t*
**while** convergence criteria **do
for each** ensemble *µ ∈* [1, *A*] time *k ∈* [1, *M*] **do**
draw *ω*_*kµ*_ *∼ P (ω*_*kµ*_|*t, s*)
**for each** cell *i∈*[1, *N*] **do**
draw *µ*^*∗*^*∼q*(*µ*)
**if** *µ*^*∗*^ = *A* + 1 **then**
draw *ω*_*kµ*_*∗ ∼ P (ω*)
*A → A* + 1
Accept new *µ*^*∗*^ with probability *a*(*µ*^*∗*^, *µ*_0_)
**if** *G*_*µ*0_ = 0 **then**
delete ensemble *µ*_0_
*A → A -* 1
draw *θ ∼ P (θ*|*t, ω, s*)
~~~

## Results

### Validation

To validate the method we generated random neuronal activity matrices *s* from our model by fixing coherence, noise and activity for each neuronal ensemble, then we used the Montecarlo algorithm 2 outlined in the previous section to infer parameters and latent variables. In Figure 2 we illustrate a sample of the activity matrix *s* obtained from the model with five ensembles with cells sorted by membership (Fig. 2A). At the initialization step we assigned the membership of each cell uniformly between 0 to *A*^*max*^ where *A*^*max*^*≈N*. In Fig. 2C we display the course of the reassignment for each neuron at four different stages of the sampling. In particular, the early phase where initial groups are formed is followed by the merging of equivalent ensembles which eventually converge to the true assigned membership. Fig. 2B displays the convergence of the log-likelihood (top panel) and the membership transition rate, defined as the number of cells reassigned to a different ensemble divided by the total number of cells (bottom panel). When the data display a strong organization in terms of synchronous activity, deviations from the maximum likelihood configuration are suppressed. Therefore once cells are assigned to their ground truth label (up to permutation) they are unlikely to be reassigned after the sampler has reached convergence.

**Figure 2:**
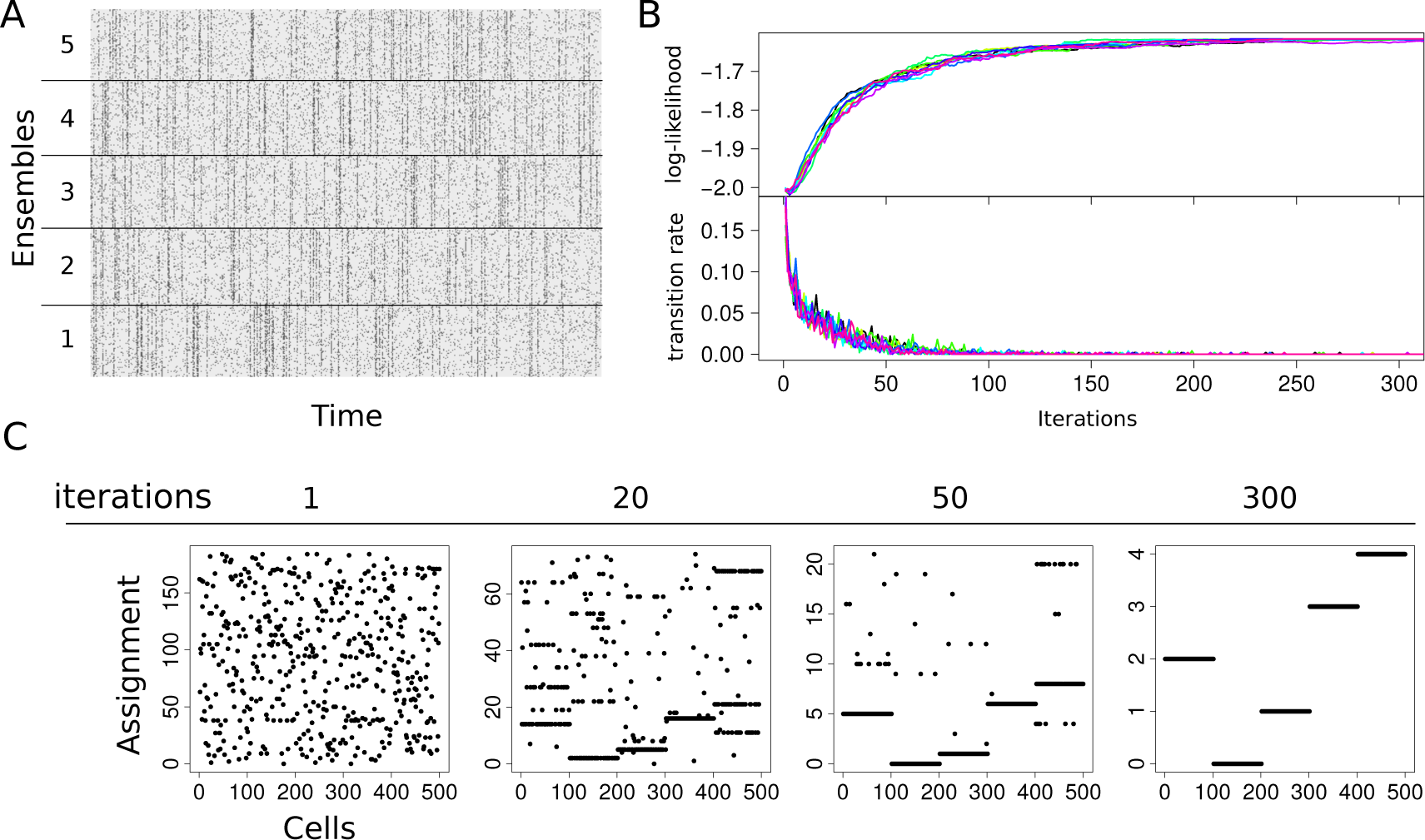
Validation. (A) Simulated activity of *N* = 500 neurons organized in *A* = 5 ensembles. Time sequences of length *M* = 1000 for all neurons have been generated from the model with *λ*_*µ*_(0) = 0.08, *λ*_*µ*_(1) = 0.6 and *p*_*µ*_ = 0.1 for noise, coherence and activity equal for all ensembles. We used this activity matrix as input to our inference algorithm to recover the five ensembles. (B) (top) log-likelihood over the course of 300 iterations of the Montecarlo sampler and (bottom) number of transitions per iteration divided by the number of neurons *N* (different colors correspond to different initializations of neuronal memberships). (C) Assignments of cells to ensembles at four stages of the sampling. Cells initially assigned to *A≈N* ensembles are progressively grouped until the original ensembles are recovered.

Next, we explored a broad range of noise and coherence to describe the regime of applicability of our method. We performed inference on random activity matrices generated from the model with noise and coherence in the range between 0.05 and 0.95. The diagram depicted in Figure 3A shows the separation between regions where all cell memberships are exactly recovered (detectable phase, red) from regions where full recovery of cell membership is not achievable (non-detectable phase, blue). In particular, ensemble recovery is possible whenever the absolute difference between noise and coherence |*λ*(1)*-λ*(0)|is sufficiently high. Indeed, the condition *λ*(1) = *λ*(0) corresponds to the degenerate case where the firing probability is independent of the ensemble state, in which case *s* does not provide any statistical information about cell membership. The width of the non-detectable region depends on ensemble activity *p*_*µ*_’s; at fixed time *M* higher ensemble activity provides more information. Therefore, when the activity increases the size of the non-detectable region is reduced (Fig. S1). The detectable phase is separated in two symmetric regions. *λ*(1) *> λ*(0) corresponds to “on” ensembles where neuron activity is correlated with ensemble activity. The detectable region with *λ*(1) *< λ*(0) corresponds to “off” ensembles where member neurons are anti-correlated with ensemble activity. This regime describes the scenario where a population of highly active neurons is characterized by synchronous transitions to “off states”. The average membership transition rate can be viewed as an order parameter for the detectable/non-detectable phase transition. In fact, unlike the detectable phase where the membership transition rate vanishes, in the non-detectable phase this quantity reaches a non-zero limit, as shown in Figure 3B, indicating that maximum-likelihood fluctuations are no longer suppressed. In this regime, the information provided by the activity matrix *s* is not sufficient to assign neurons to an ensemble.

**Figure 3:**
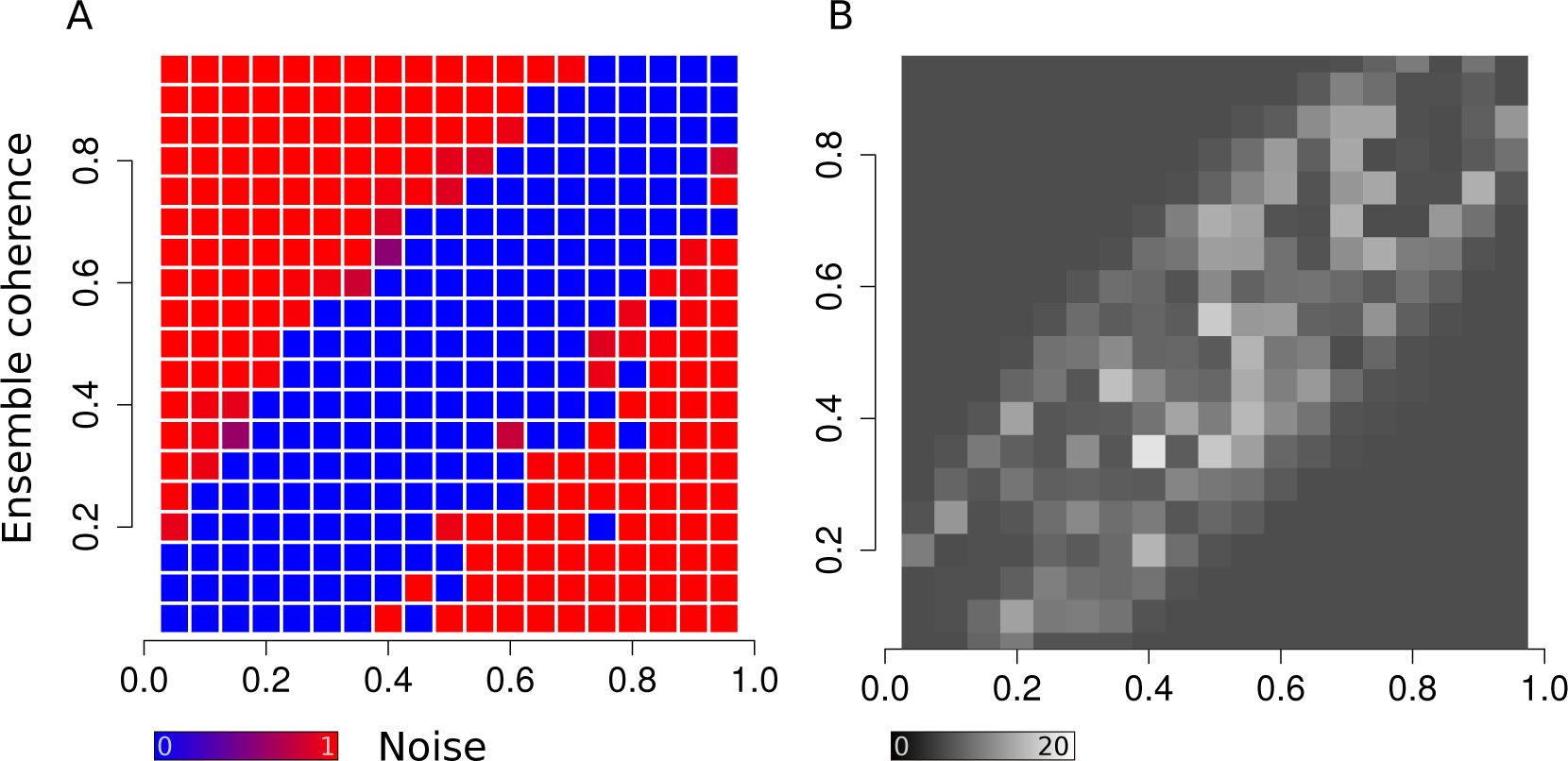
Phase diagram. (A) Raster plot displaying detectable (red) versus non-detectable (blue) inference of 5 ensembles randomly generated using different values of noise and coherence. In particular ensemble inference is non-detectable when noise and coherence are numerically close. (B) Raster plot representing the number of neurons reassigned, on average, for every random sample of the Markov chain. The average transition rate can be viewed as an order parameter characterizing the transition between detectable and non-detectable phases.

### Spontaneous activity in the zebrafish tectum

We used our method to infer the precise structure and dynamics of spontaneously active neural ensembles within the optic tectum of the larval zebrafish. For these experiments we used zebrafish with pan-neuronal expression of a nuclear localized calcium indicator, Tg(*elavl3:H2B-GCaMP6s*) (see the Methods section). Using two-photon volumetric calcium imaging, we monitored spontaneous activity in the tectum of immobilized zebrafish larvae kept in total darkness. In all experiments we recorded calcium activity for 1 hour throughout 5 planes, 15*µ*m apart with an acquisition frequency of 4.8Hz per volume (Fig. 4A, Movie S1 and the Methods section). Volumetric images were preprocessed using registration and segmentation software to extract the time sequences from thousands of tectal cells in both tectal hemispheres (Fig. 4B and see Methods section).

**Figure 4:**
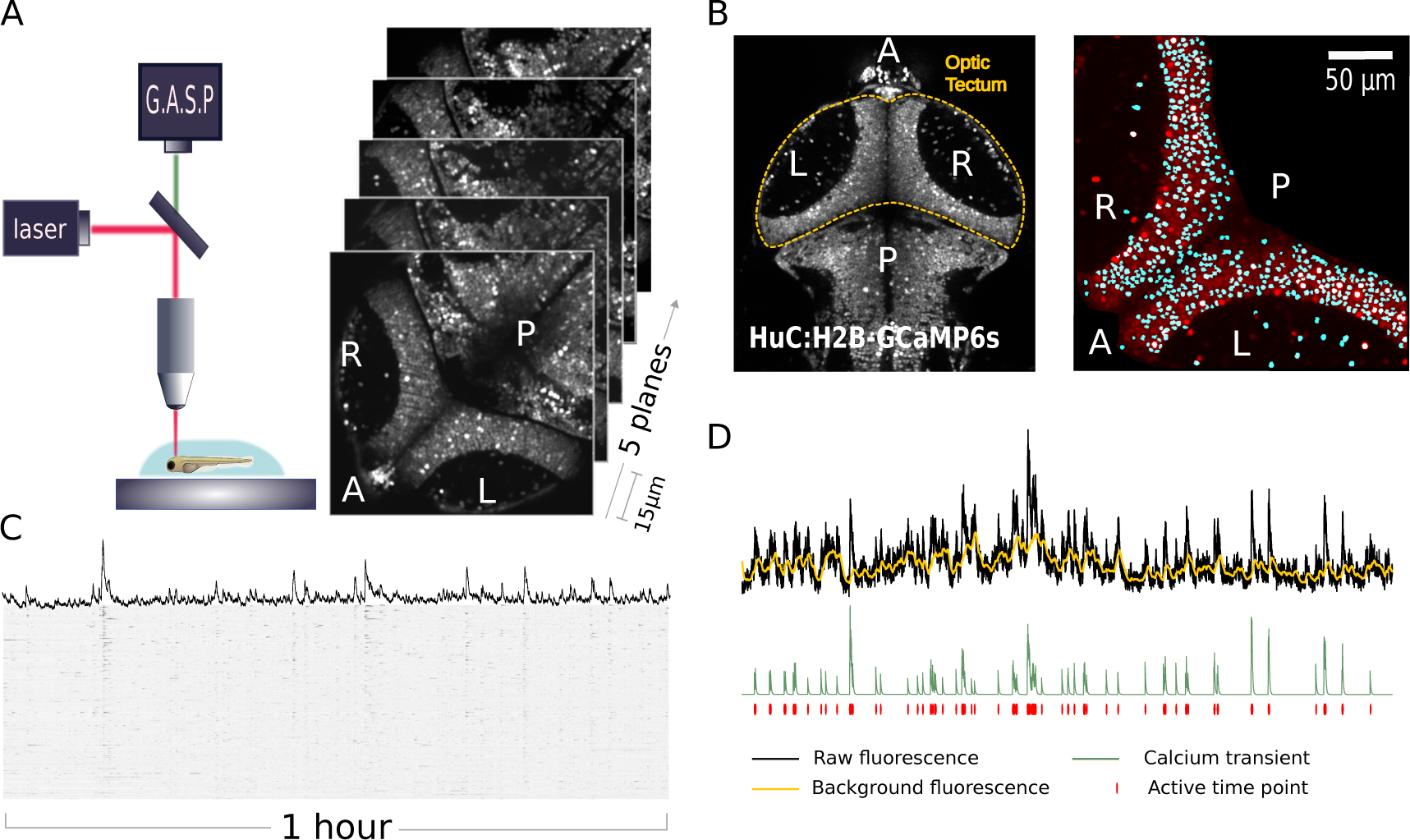
Imaging of the zebrafish tectum. (A) Volumetric 2-photon imaging of both tectal hemispheres. System setup (left) and time averages of raw fluorescence showing anterior/posterior axis (A/P) and right/left (R/L) optic tectum (right) (see also Movie S1). (B) Image processing and cell segmentation. (C) Raster plot showing calcium activity over the course of the experiment. (D) Hidden Markov model for extracting the calcium transient times.

In order to match the data with our generative model we extracted the onset times of each calcium transient from all tectal cells in the form of a binary matrix (Fig. 4C). To do so we separated calcium transient signals from baseline activity by using a hidden Markov model (HMM) where the neuronal activity (hidden) state *s*_*t*_ at time *t* is represented by a Bernoulli process, the background *b*_*t*_ is a Gaussian Markov process and the calcium transients are drawn from a normal distribution when *s*_*t*_ = 1 or follow a deterministic exponential decay when *s*_*t*_ = 0 (Fig. 4D). More realistic models have been developed to describe the fluorescence levels of the calcium indicator[10]. However, since our purpose is to capture the synchronous events among cells and not to infer the number of spikes within a calcium transient we can use a simplified model to select events where the activity deviates significantly from the background. The HMM approach can be used to obtain analytical estimates of signal and background (see Methods section) allowing fast preprocessing the calcium traces.

Estimates of the binary activity for each neuron were then combined into matrices of dimension [*N* cells] *×*[*M* time frames] where *M* = 17460 frames for all experiments and a number of neurons *N* of order 10^3^ dependent on how many cells were segmented for each fish. To build the activity matrix *s* used for inference we considered all time frames with more than 15 active cells. We applied our method to each binary activity matrix by running the inference algorithm 2 until convergence. The advantage of our method is that ensemble features (activity, noise and coherence) are simultaneously estimated together with neuronal membership, which allows us to select for further analysis ensembles with higher biological relevance. Ensembles with low activity levels for instance, which are composed of cells which were almost never active during the recording, can be discarded. Similarly, groups of poorly correlated neurons incoherently firing can be combined into ensembles where coherence and noise are very similar, therefore potentially outside the detectable regime. This allows us to automatically separate “free” neurons, which do not belong to a coherent ensembles, from “ensembled” neurons where the coherence is significantly larger than the noise level. To focus on the coherent populations we selected ensembles with activity larger than 0.5% (corresponding to*≈*1 event per minute), size larger than 5 neurons, coherence larger than 5% and noise lower than 5%.

By comparing the analysis of *n* = 7 fish using our technique, we obtained a number of neuronal ensembles (Fig. 5A) which varied between 30 and 50 per larva. All ensembles features displayed non-significant variations across fish (Fig. 5B). In particular, the average ensemble coherence across fish was 0.19 (SD = 0.11), corresponding on average to a 19% chance for a member neuron to be recruited during ensemble activation. In addition, we have found typical noise levels of 0.007 (SD = 0.005), meaning that there is a 0.7% chance of a neuron to be active at any time point where its ensemble is off. Although the average coherence seems low, this is because our definition of coherence is a measure of synchrony at equal times (within the temporal resolution of the recording). However, ensemble activity is characterized by more sustained periods of activation during which most member neurons are recruited gradually rather than instantaneously (see Fig. S4). However our inference method softens this constraint by taking into account sustained activity by extending the time window in which an ensemble is active. This allows us to accommodate ensembles where synchrony occurs over extended periods of time rather than single time points.

**Figure 5:**
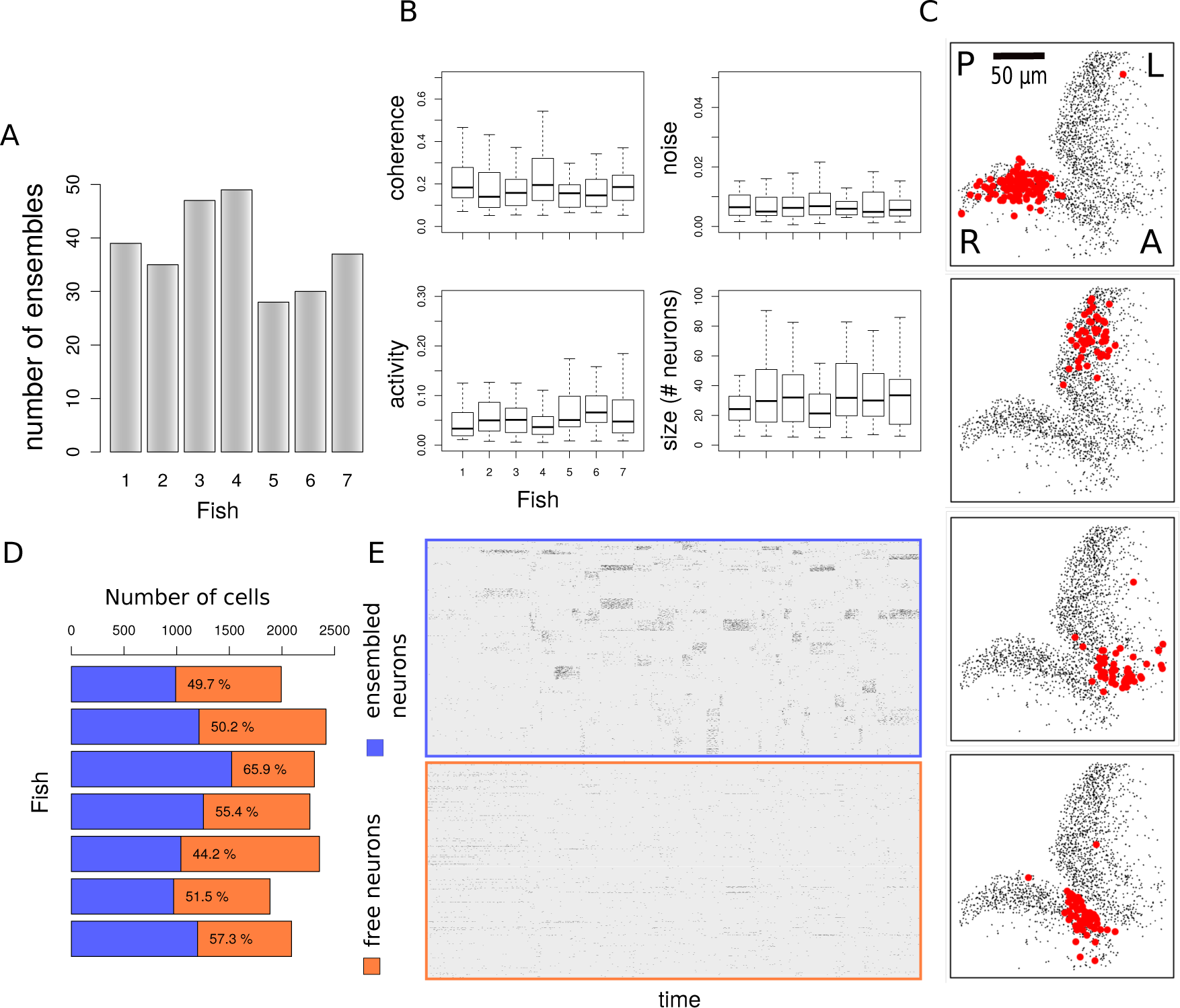
Stereotyped features of neuronal ensembles across fish. (A) Number of neuronal ensembles estimated for each fish. (B) Comparison of coherence, noise, activity and ensemble sizes for all experiments. Coherence and noise correspond to the model parameters *λ*(1) and *λ*(0), as the probabilities of a neuron to fire with its ensemble (coherent activation) and independently of it (noisy activation). The activity corresponds to the probability of an ensemble to be active at any time during spontaneous activity events. (C) Representative ensembles from single fish with high coherence and activity display spatial compactness within left or right tectal hemispheres. Labels A/P and R/L indicate anterior/posterior axis and right/left optic tectum respectively. (D) Fraction of neurons assigned to ensembles with 99% confidence. The efficacy of our method in capturing neuronal ensembles is illustrated by separating the recording of ensembled neurons from free neurons (E). When sorting the activity matrix by membership, ensembled neurons generate bands of synchronous activity whereas free neurons display independent random firing events.

Next we examined how the ensembles selected according to their features are distributed in the tectum. We found that over 90% of all detected ensembles were spatially compact within one of the two tectal hemispheres (Figs. 5C, 6D) and in their 3D distribution (see Movie S2). In some of the ensembles we observed the presence of a small number of “satellite” cells which were spatially separate from the ensemble core and which were located either in the ipsilateral or contralateral tectum. Up to 66% of the spontaneously active neurons were assigned to one of the selected ensemble with 99% of confidence (Fig. 5D), showing that neuronal ensembles recruit over a half of the tectal population over 1 hour of recording (Fig. 5E).

The estimated ensemble activity over time (Fig. 6A) provides a coarse-grained description of the spontaneous activity in the optic tectum (see Movie S3). Although we did not assume any correlation among the ensembles, we can use the ensemble time sequences inferred from the data to explore their relationships in the form of network graphs. The (Pearson’s) time correlation between neuronal populations (Fig. 6B, see also the Method section) revealed the presence of subnetworks of ensembles (Fig. 6C). These subnetworks are largely composed of neighboring ipsilateral ensembles with rarer edges between ensembles in opposite tectal hemispheres. This is shown in Fig. 6C by representing the Pearson’s correlation matrix between ensemble time sequences as a graph with nodes corresponding to the neuronal ensembles and edges representing a positive correlation (with 95% confidence). Shorter edges reflect higher time correlation. Inspection of these graphs reveals that edges between ensembles located in anterior and posterior regions of a tectal hemisphere are almost never observed, as shown by the decrease of time correlation at large spatial separation between ipsilateral pairs of ensembles (Fig. 6E-F). These results suggest functional segregation of anterior and posterior tectal networks. Moreover, the degree of correlation between ensembles located in opposite tectal hemispheres (even at a similar topographic location) is generally lower than the correlation between ipsilateral ensembles.

**Figure 6:**
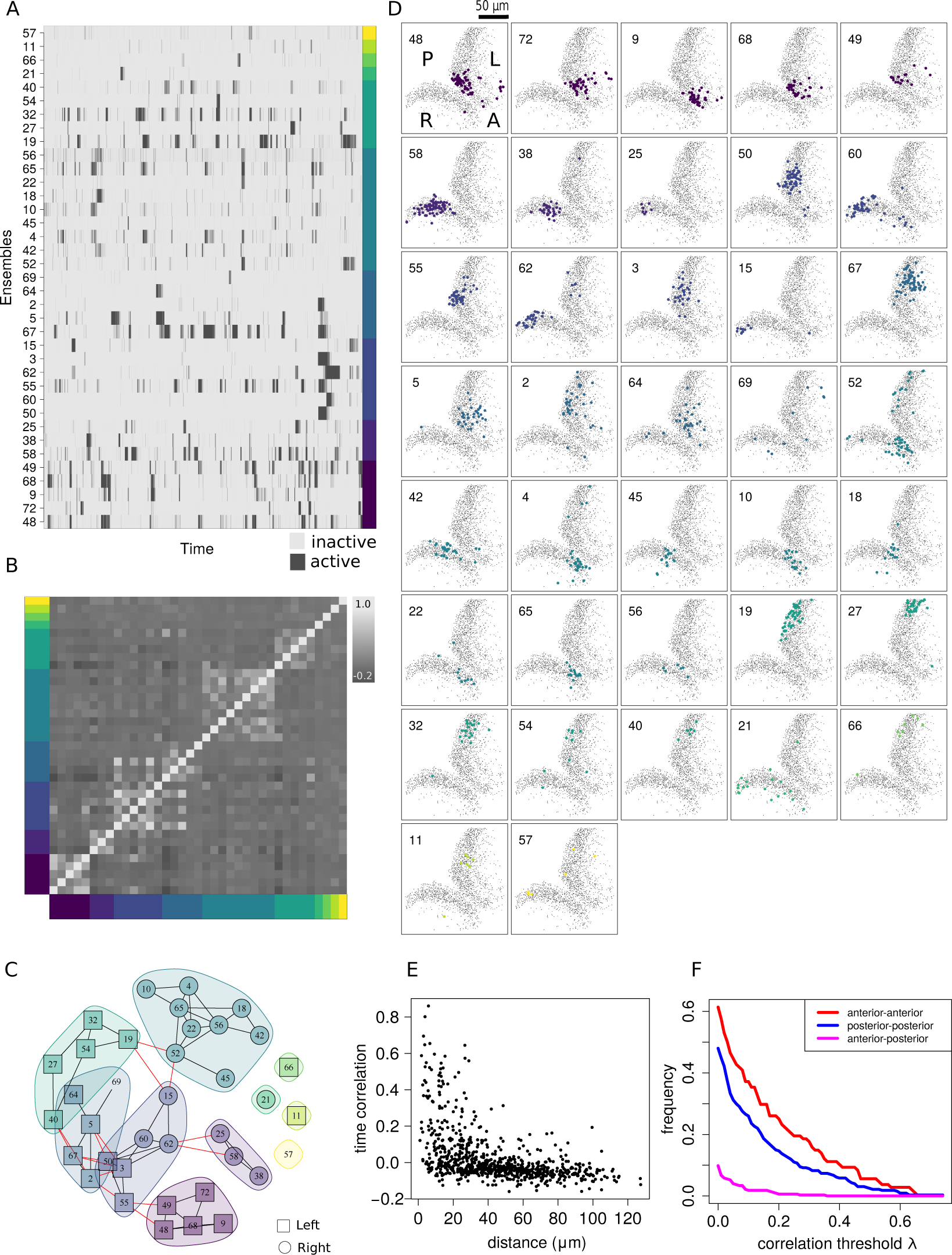
Networks of ensembles in the zebrafish tectum. (A) Raster plot of ensemble activity ordered according to subnetwork membership as shown in C. (B) Raster plot of the time correlation matrix (Pearson’s) between all pairs of ensembles averaged across posterior samples and sorted according to subnetwork membership. (C) Subnetwork graphs representing the time correlations among neuronal ensembles (see the Method section). Each ensemble is a node, shorter edges reflect higher temporal correlations. Note the absence of edges between ensembles located in ipsilateral anterior and posterior tectum (see Movie S3). (D) Locations of all ensembles from a single larva (see also Fig. S3 for a different larva and Movie S2 for the ensemble distributions in 3D). Each ensemble is color-coded according to subnetwork memberships shown in C. Neighboring ensembles form subnetworks. (E) Scatter plot showing the relationship between time correlation and physical distance between neuronal ensembles. (F) Fraction of ensemble pairs with correlation larger than *λ* as a function of *λ*. Ipsilateral pairs of ensembles located at opposite sides in the anterior-posterior axis display a significant reduction in correlation compared to neighboring ensemble pairs (anterior-anterior and posterior-posterior).

### Analysis of functional imaging data from the mouse cortex

To illustrate the general applicability of our method we carried out analysis of neuronal ensembles in the mouse cortex (Fig. 7A). We analyzed a dataset generated by Stringer et al.[34, 33] where the activity of over 10,000 neurons in the mouse visual cortex were recorded in the absence of external stimulation but head-fixed mice were free to run on a air-floating ball. As we did for the zebrafish tectal data, we first applied our HMM technique (see Methods) to extract calcium transients from the raw fluorescence of each recorded neuron to obtain the binarized neuronal activity matrix. Then we applied our bayesian model to detect neuronal ensembles from the binary activity and infer their properties. Figure 7B displays the ensemble activity over time for all ensembles satisfying the same selection criteria of activity, size, coherence and noise used for the zebrafish tectum. The majority of the cortical ensembles are distributed across the volumed imaged (see Movie S4a representing ensemble 24), with few exceptions with neurons organized into columns spanning through the z-axis (Movie S4b, ensemble 43) or limited to ventral or dorsal planes (Movie S4c, ensemble 23). We analyzed the time correlation between ensemble activities obtained with our method and some of the behavioral features (running speed and pupil area) which were simultaneously monitored during functional imaging in Ref. [[34]]. Several ensembles are characterized by a remarkably positive or negative correlation with running speed, consistent with the finding in Ref. [[34]] that the first principal component of the neuronal activity is highly correlated with arousal. In particular, the activity of the most populated ensemble (ensemble 24, Fig. 7C) is anti-correlated with running speed, as opposed to ensemble 15 shown in Fig. 7C-D which tends to be active during periods of running. Ensembles can be correlated with running speed at different time points or differ in activity and coherence (see Fig. S6 for a comparison between ensembles 15 and 32), suggesting that arousal is associated with the coordinated activation of multiple ensembles. By using the time course of the pupil area we constructed a binary vector indicating the events of pronounced pupil contraction. One of the detected ensembles (ensemble 13, Fig. 7C) is highly correlated with pupil contraction events, as shown in Fig. 7E.

**Figure 7:**
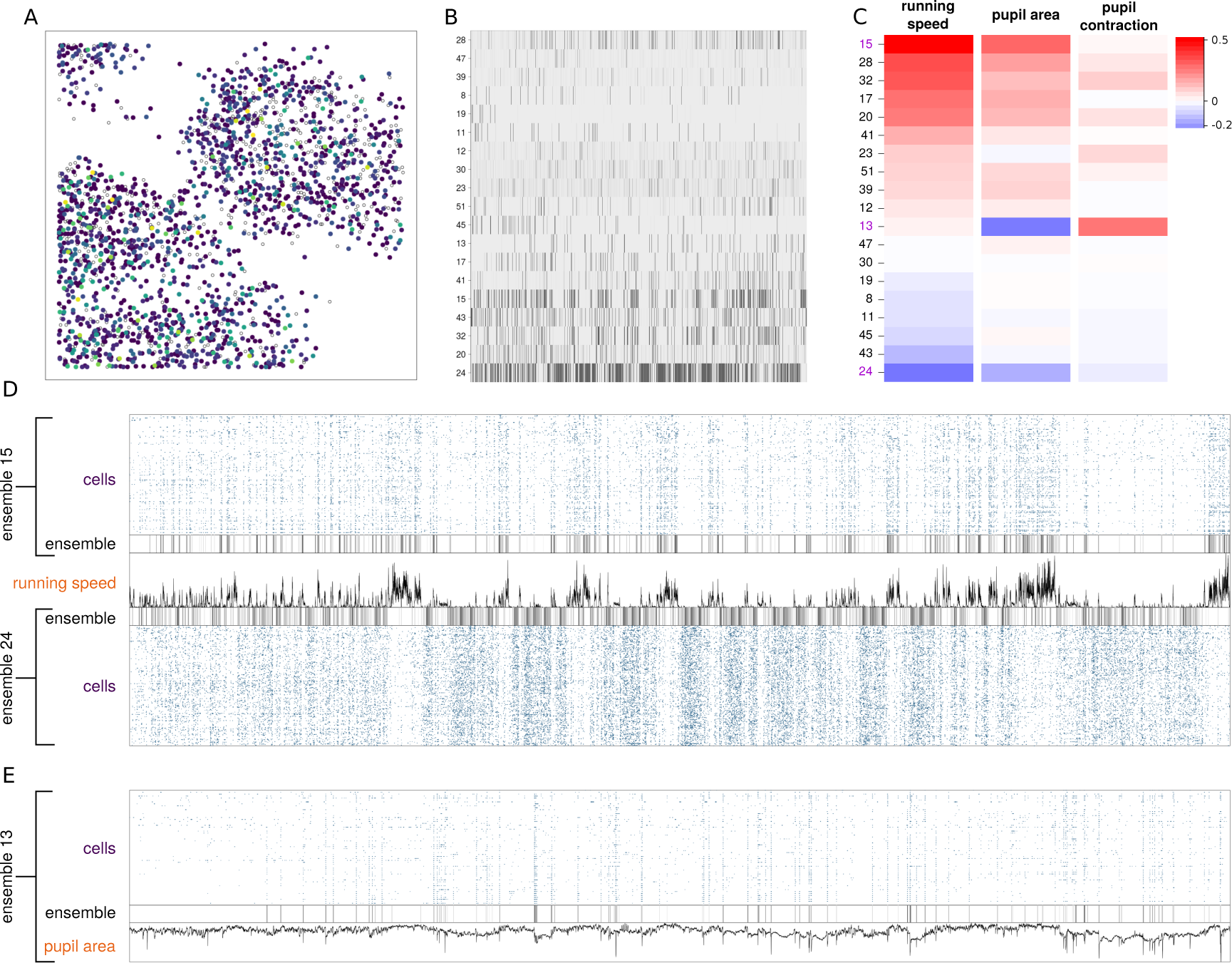
Neuronal ensembles from functional imaging of the mouse cortex. (A) XY distribution of recorded neurons[34] color coded according to ensemble membership. (B) Ensemble activity matrix. (C) Correlation between the activity of each ensemble and (from left to right) the time course of running speed, pupil area and the (binarized) pupil contraction. (D) Correlation and anti-correlation of neuronal ensembles with respect to running speed. (E) Ensemble correlated with pupil contraction.

### Identification of ensembles in neuropixels recordings in mouse

Neuropixels electrode arrays permit large-scale electrophysiology by recording from hundreds of neurons simultaneously. We have analyzed the data from Ref. [[32, 23]] where a neuropixels probe was used to record the activity from the visual cortex, hippocampus and parts of the thalamus in a head-fixed mouse in the absence of stimulation with forepaws resting on a wheel that could move laterally. The dataset contains 242 isolated neurons for which we analyzed the ensemble structure using our bayesian method.

In order to carry out the analysis of the ensembles from large-scale electrophysiological recordings, we need the neuronal activity matrix as input of our inference method. Although it would be natural to define the state of a neuron according to the presence or absence of a spike at any given time point, this definition is not suitable in this context because the synchronous activation of a neuronal population occurs at a much slower time scale than a single spike. Here we define the state of a neuron based on the firing rate instead of single spikes. Periods of increased firing rate due to reverberating activity of recurrent networks[20] can indeed last for tens of seconds, matching the time scale of ensemble activity.

We extracted the transients of high firing frequency for each neuron by counting the spikes in bins of 0.6s along the recording and then applying the HMM method to detect the transient onsets (Fig. 8A). By applying our inference method to the binary activity matrix representing the states of all 242 neurons over time (Fig. 8B), we found that 33% of those neurons can be assigned with 99% confidence to four ensembles satisfying the same constraints on activity, coherence and noise as applied to calcium imaging data). In particular ensembles 1 and 3 displayed a significantly positive correlation with wheel velocity (reflecting lateral mouse movement) and ensemble 2 was weakly anti-correlated with wheel velocity (Fig. 8E-F). The observation of neuronal ensembles either positively or negatively correlated to the motor output is consistent with the analogous results obtained by analyzing functional imaging data of the mouse cortex. These results and those obtained from the functional imaging data from mouse and zebrafish demonstrate that our method is able to identify ensembles in diverse species and data types.

**Figure 8:**
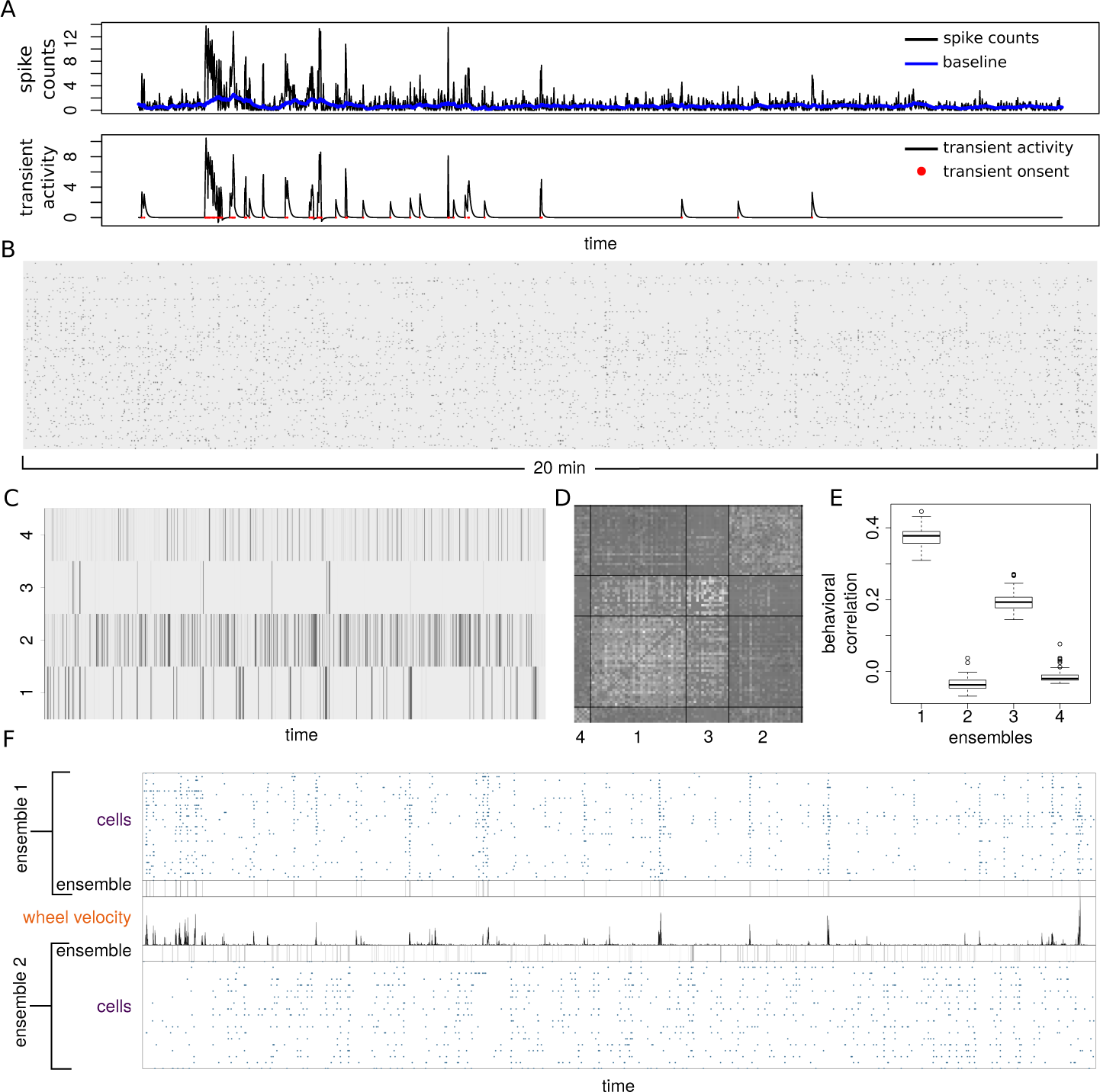
Ensembles from neuropixels recording of mouse cortex. (A) Identification of activity transients with increased firing frequency from spike counts using bin size of 0.6s. (B) Binary activity matrix obtained by combining transient events from 242 neurons. (C) Raster plot of the ensemble activity. (D) Raster plot of the time correlation matrix between neurons sorted by ensemble membership. (E) Time correlation between each ensemble activity and the absolute wheel velocity. Boxplots represent the distribution of correlation across posterior samples. (F) Comparison between correlated and anti-correlated ensembles with respect to wheel motion.

## Discussion

In this work we introduced a novel, model-based approach to detect neuronal ensembles from spontaneous activity. Our method provides a full characterization of these neuronal ensembles by simultaneously quantifying their composition, noise level, activity rate and within-ensemble synchrony. Furthermore, it generates a coarse description of the neuronal dynamics by inferring the activation states of each neuronal population over time, which can then be used to uncover relationships between ensembles. Within our model we introduced three key and novel features to characterize ensembles: coherence, which is a measure of the level of synchrony among neurons within an ensemble; noise, which quantifies the probability of the member neurons to fire when the ensemble is inactive, and ensemble activity which is the probability of the ensemble to be active.

By applying our method to functional imaging data from the zebrafish optic tectum, we revealed a high level of biological reproducibility when comparing ensemble features across fish. An additional finding is that the majority of the ensembles are spatially compact and can be grouped into segregated subnetworks restricted to either the anterior or posterior half of a tectal hemisphere. These findings are consistent with optogenetic data showing that stimulation of anterior or posterior tectum drives different (approach- and avoidance-like) behaviors. Therefore, the functional segregation of subnetworks of ensembles may be relevant for understanding how activation of different tectal locations leads to mutually exclusive behaviors[17, 12, 31]. Due to our simultaneous analysis of ensemble structure and features we could focus on the neuronal populations which displayed biologically relevant activity and coherence, revealing that a large fraction of the tectal neurons could not be assigned into a specific neuronal ensemble. These “free” neurons may be part of ensembles which were inactive during our recording period. This may be due to chance or there may be some tectal ensembles which are only recruited during sensory stimulation and/or behavior. Alternatively, some of the free neurons might not belong exclusively to one ensemble. Although our generative model assumes that each neuron belongs exclusively to one ensemble, the multimodality in the posterior membership distributions may reflect the presence of such “promiscuous” neurons. To accommodate soft assignment it is possible to extend the model by introducing latent variables describing the probabilities of each neuron to be recruited by any of the ensembles. This extension however might require further development of the Markov chain Montecarlo algorithm in order to sample efficiently from the posterior distribution. A final interpretation is that the free neurons are only weakly coupled to population firing. Such neurons, referred to as “soloists”, have been identified in the mouse and macaque visual cortex where they contrast with “choristers” whose firing is strongly coupled to the overall firing of the population[28].

We illustrated the general applicability of our inference method by also analysing the neuronal ensembles present in shared functional imaging data and neuropixels recording from the mouse. By exploiting the behavioral information recorded alongside neuronal activity, we have found that some of the ensembles detected with our method were correlated or anti-correlated with specific motor outputs such as running or pupil contraction. These findings agree well with those of the Stringer et al[34, 33], who show using principal component analysis that the first principal component correlates with arousal (as indicated by running speed). Our method builds on these findings by revealing the specific neural ensembles that differentially correlate with arousal at different time points or with different levels of activity and coherence. Our method provides a statistically rigorous means to interrogate the differences between such ensembles further, for example by quantifying differences in coherence, noise, activity, size and functional interactions with other ensembles.

In this work we demonstrated the applicability of our framework in the context of calcium imaging and neuropixels recording data by analyzing time sequences of thousands of neurons to detect synchronously active populations. By extracting key ensemble features from the data we have provided new insights into the spatiotemporal organization of the zebrafish tectum and link ensemble dynamics in the mouse cortex to behavioral output. More generally, our approach may be used to quantify how ensemble structure and dynamics are remodeled as a function of development, aging, experience, internal states and disease. From a statistical perspective, our inference method is not limited to neuronal population activity data and could be applied to any dataset where a large number of agents perform actions over a long period of time to reveal the groups of agents displaying coordinated behavior.

## Methods

### Zebrafish larvae

For functional imaging experiments we used transgenic zebrafish, Tg(*elavl3:H2B-GCaMP6s*), expressing the nucleus-targeted calcium indicator GCaMP6 (Chen et al., 2013) under the *elavl3* promoter, which provides near-panneuronal expression (gift from Misha Ahrens, Janelia Research Campus). Larvae were raised at 28.5^°^C in Danieau solution and were exposed to a 14 hour ON/10 hour OFF light/dark cycle. Larvae were fed daily from 5 dpf using live rotifiers. To maximize optical clarity for imaging Tg(*elavl3:H2B-GCaMP6s*) larvae were crossed with compound *roy;nacre* double homozygous mutants (*casper*) larvae which lack melanocyte and iridophore pigmentation. This work was approved by the local Animal Care and Use Committee (Kings College London), and was carried out in accordance with the Animals (Experimental Procedures) Act, 1986, under license from the United Kingdom Home Office.

### Volumetric Calcium Imaging

At 7 days post fertilization (dpf) zebrafish were mounted in 2% low melting point agarose in Danieau water with their dorsal side facing up on a custom built imaging slide and were submerged in Danieau water. These fish were left for 1 hour in the light so that the fish could settle, reducing drift while imaging. Prior to imaging the fish were placed under the microscope objective in total darkness for 30 minutes to allow the fish to adjust to the imaging conditions. Spontaneous activity was monitored by imaging the calcium dynamics of thousands of neurons in both tectal hemispheres for 1 hour with a custom built 2-photon microscope (Independent NeuroScience Services, INSS). Excitation was provided by a Mai Tai HP ultrafast Ti:Sapphire laser (Spectraphysics) tuned to 940nm. Laser power at the objective was kept below 15 mW for all fish. Emitted light was collected by a 16x, 1 NA water immersion objective (Nikon) and detected using a gallium arsenide phosphide (GaAsP) detector (ThorLabs). Images (256 × 256 pixels) were acquired at a frame rate of 60Hz by scanning the laser in the x-axis with a resonant scanner and in the y-axis by a galvo-mirror. To enhance signal-to-noise every 2 frames for each focal plane were averaged. The focal plane was adjusted in 15*µ*m steps using a piezo lens holder (Physik Instrumente). This allowed for volumetric data consisting of 5 focal planes to be collected at a volume rate of 4.8Hz. Scanning and image acquisition were controlled by Scanimage Software (Vidrio Technologies).

### Image registration

Any volumetric stacks where large numbers of neurons in the imaging plane drifted out of view were discarded. All other stacks were corrected for x-y drift by aligning each every frame in each slice in the stack independently. First every frame for each slice was aligned to its first frame, then they were aligned again to the mean of these images. This registration was performed using a non-rigid body alignment algorithm contained within the SPM8 package for MATLAB (http://www.fil.ion.ucl.ac.uk/spm/software/spm8).

### Segmentation of tectal cells

After image registration, images were processed using custom-made C++ software of cell segmentation based on a flooding algorithm. For each plane in the volume, we generate the time average of the image and apply a Gaussian filter to smooth the spatial details below the size of a cell. Next, we applied the following segmentation algorithm to extract single cell time sequences. Starting from the top fluorescence *f* ^(*max*)^ we sequentially label voxels according to the nearest segmented cell within a radius compatible with the size of a cell. To take time correlations into account we introduced a distance between voxels *h*(*V, V* ^*’*^) which combines physical separation and time correlation as

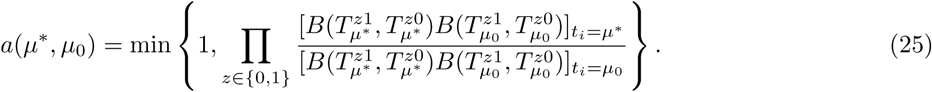

where the parameter *η* introduce some flexibility in the importance of the Pearson’s time correlation *C*_*V* *V*_′ between the time sequences of *V* and *V′* Whenever a new label is assigned, the corresponding voxel becomes the representative of the new cell and the distance between a voxel *V* and a cell is obtained by Eq. (26) by replacing *V’* with the voxel which representing the cell.

#### Algorithm 3 Segmentation algorithm

~~~
1: Set *f* = *f* ^(*max*)^
2: **while** *f ≥ f* ^(*min*)^ **do**
3: **for each** voxel *V* with fluorescence *≥ f* **do**
4: Calculate the distance *h* between *V* and its nearest cell according to Eq. (26)
5: **if** *h < h*^(*max*)^ **then**
6: Assign *V* to the nearest cell
7: **else**
8: Assign *V* to a new label
9: *f* = *f -*Δ*f*
~~~

### Detection of calcium transients

As discussed in the main text, we decompose fluorescence traces 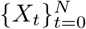in the sum of calcium transient *c*_*t*_, baseline activity *b*_*t*_ and a source of Gaussian noise. The hidden Markov model is summarized by the set of equations

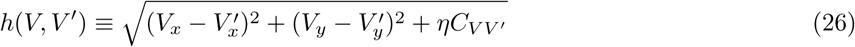

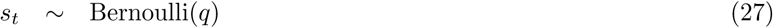

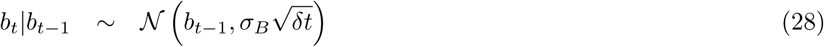

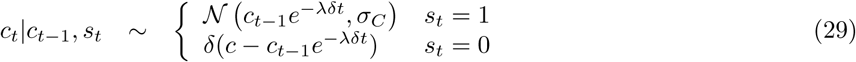

where calcium transients are specified by the latent variable *s*_*t*_ representing the hidden on/off state of a neuron, modeled as a Bernoulli process. To obtain an estimate of the hidden variables at all times we maximize iteratively the probability of the latent state at time *t*, given the state at *t -* 1 and the observed value *X*_*t*_

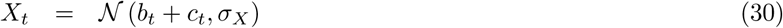

where all the probabilities can be obtained from the definition of the model

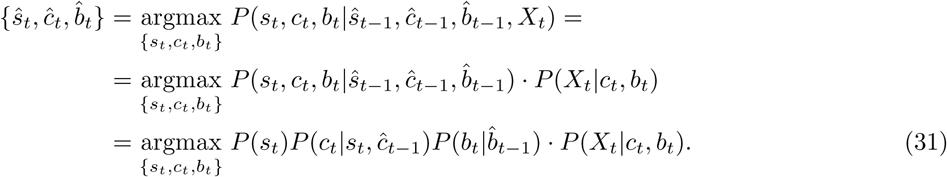

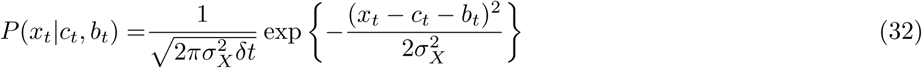

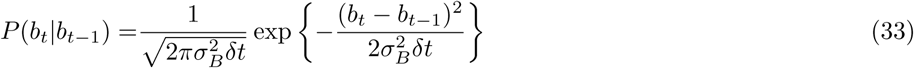

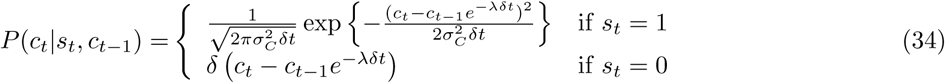

We can obtain analytical expression for the estimated calcium and baseline at each time by considering separately the two cases *s*_*t*_ = 0 or *s*_*t*_ = 1.

- **case** *s*_*t*_ = 0. To maximize the probability we need to maximize the function

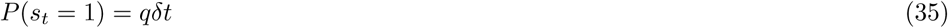

where we defined 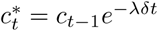. By setting 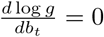 we get

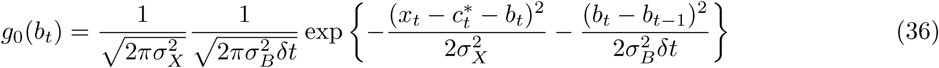

therefore the values of *c*_*t*_ and *b*_*t*_ which maximize the log-likelihood when *s*_*t*_ = 0 are

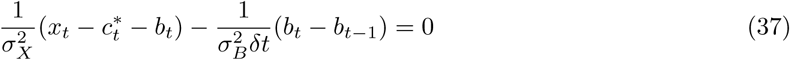

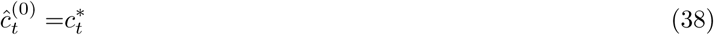
- **case** *s*_*t*_ = 1. In the case of a calcium transient we need to maximize the function

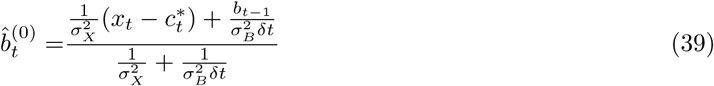

analogously to the case *s*_*t*_ = 0, by maximizing log *g*_1_ with respect to both *c*_*t*_ and *b*_*t*_ to zero we get

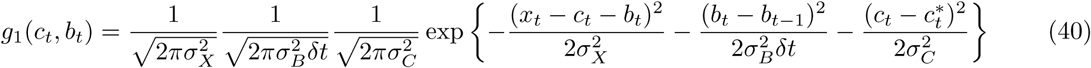

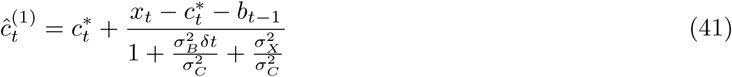

By combining the two conditions we have

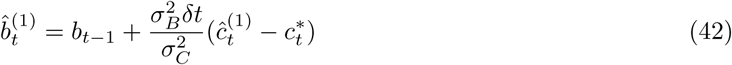

To complete the iterative algorithm we have to assign values to the initial calcium and baseline level. A simple choice for the initial state is *b*_0_ = *X*_0_ and *c*_0_, *s*_0_ = 0 however other criteria may be more effective if transients are expected to occur at the beginning of the recording.

This procedure can be applied to any time sequence to estimate the onset of exponentially decaying transient, however it requires the variance parameters *σ*_*X*_, *σ*_*B*_ and *σ*_*C*_ as well as the decay constant *λ*. Furthermore, using the same set of parameters for all traces might lead to estimation biases. To account for parameter variation across cells we employed a “plug-in” refinement method similar to the *k*-means algorithm. Starting from an initial guess of the parameters we apply the two steps

1. Generate estimats of {*s*_*t*_, *c*_*t*_, *b*_*t*_} using Eq. (43)
2. Use {*s*_*t*_, *c*_*t*_, *b*_*t*_} to obtain new parameters according to

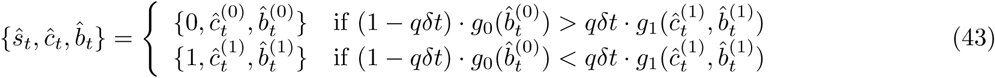

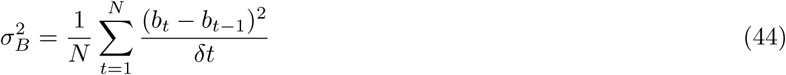

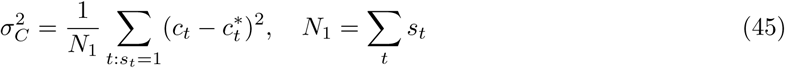

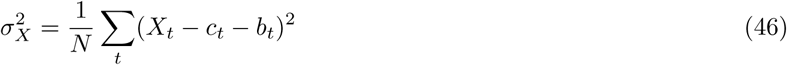

Because calcium and baseline estimated according to Eq. (43) are an approximation of the true maximumlikelihood trajectory, repeating these two steps might lead to singular values of the parameters. The first iterations of this plug-in method however provide a better decomposition than using fixed parameters. The binary activity matrices used as input for our inference method were generated by iterating twice the steps 1 and 2.

### Graph theoretic analysis of Pearson’s correlations

To carry out the time correlation analysis between ensembles in the fish tectum, for each posterior sample of the matrix *ω* representing the activity of each ensemble by row, we calculated the Pearson’s correlation between all rows. The graph describing the ensemble interaction was constructed by adding an edge between all pair of ensembles which had a positive correlation in over 95% of the posterior samples. The Girvan-Newman algorithm[27] implemented in the igraph[9] R package (edge.betweenness.community function) was used to identify the components of the correlation graph.

## Supporting information

## Acknowledgements

This work is dedicated to the memory of Diego Diana. The authors would like to acknowledge QueeLim Ch’ng and Setsuko Sahara (King’s College London) for providing HPC machines used for data analysis. Diana Passaro (Francis Crick Institute, London), Marco Banterle (London School of Hygiene and Tropical Medicine), Juan Burrone and Adil Khan (King’s College London), Marcus Triplett, Jan Moelter and Geoffrey Goodhill (Queensland Brain Institute, Brisbane) for discussions and comments on the manuscript. Misha Ahrens (HHMI Janelia) for providing Tg(*elavl3:H2B-GCaMP6s*) zebrafish. Carsen Stringer (HHMI Janelia) and Nicholas Steinmetz (University College London) for providing functional imaging data and neuropixels recordings of the mouse cortex. This work was supported by a Wellcome Trust Investigator Award MPM:204788/Z/16/Z.

## Author Contributions

G.D. and M.M. conceived the project. G.D developed the method, designed and implemented computer programs and analyzed the experimental data. T.S performed the experiments and carried out image processing and data curation. G.D., M.M. and T.S. wrote the manuscript, M.M. supervised and coordinated research activity and secured funding.

## Supplementary information

**Figure S1:**
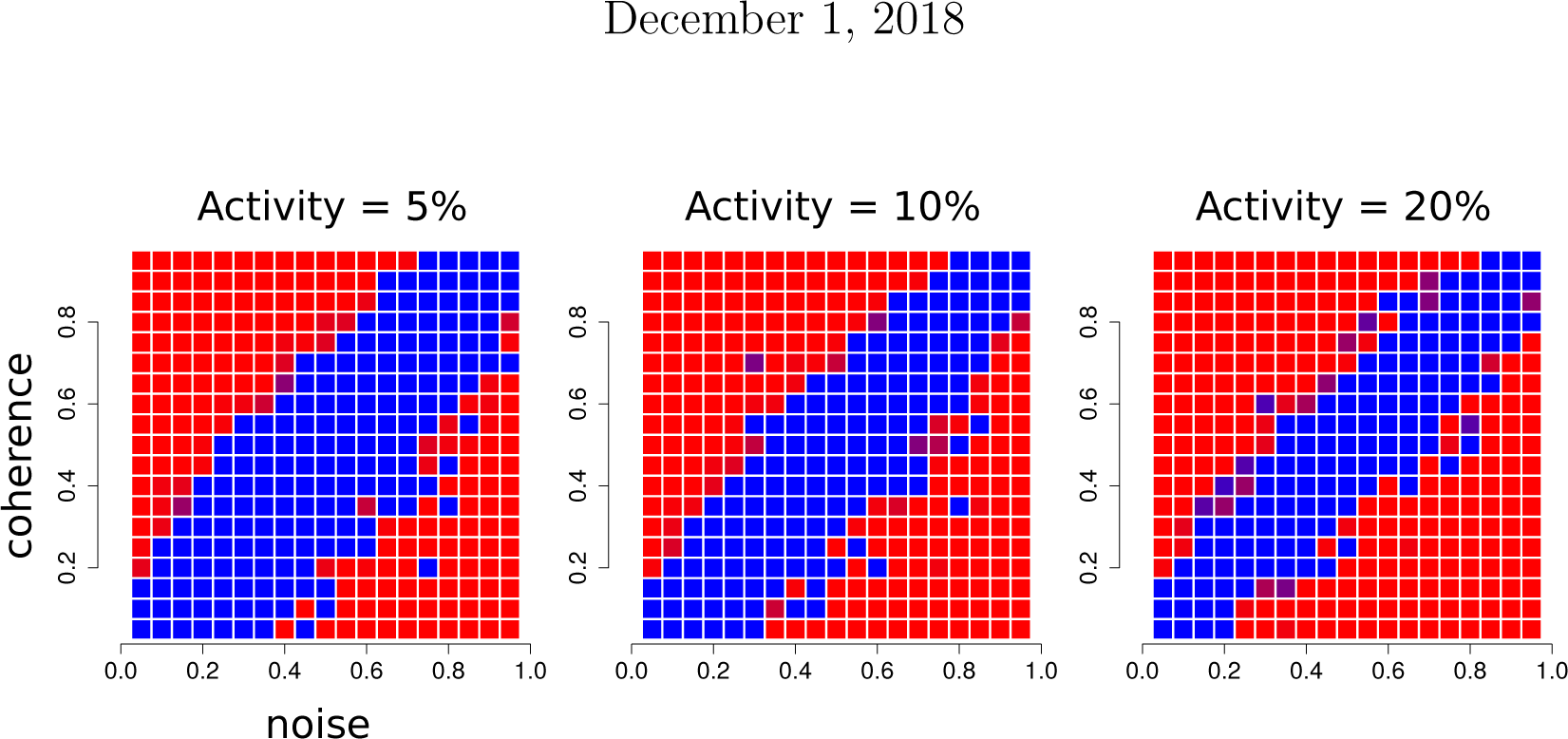
Effect of ensemble activity on the phase diagram. At increasing ensemble activity, the non-detectable regime shrinks towards the line *λ*(0) = *λ*(1), corresponding to the limit where the data are non longer informative about neuronal identity.

**Figure S2:**
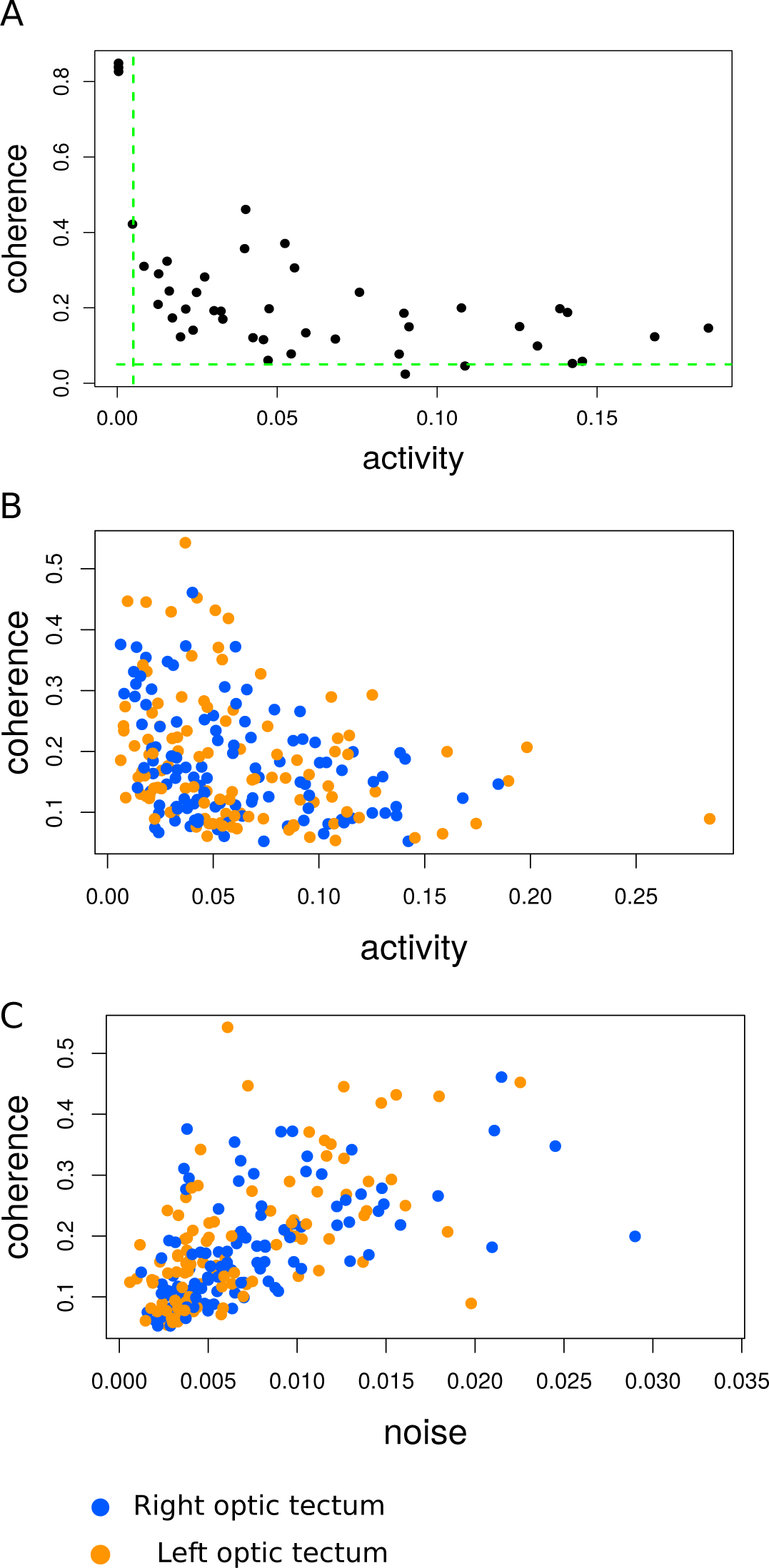
Distribution of noise, activity and coherence. (A) Scatter plot representing the distribution of noise and coherence within one fish. Green (dashed) lines display the thresholds applied to activity (0.005, corresponding to*≈*1 event per minute) and coherence (0.05). (B,C) Distribution of coherence and noise versus activity for all ensembles from all fish.

**Figure S3:**
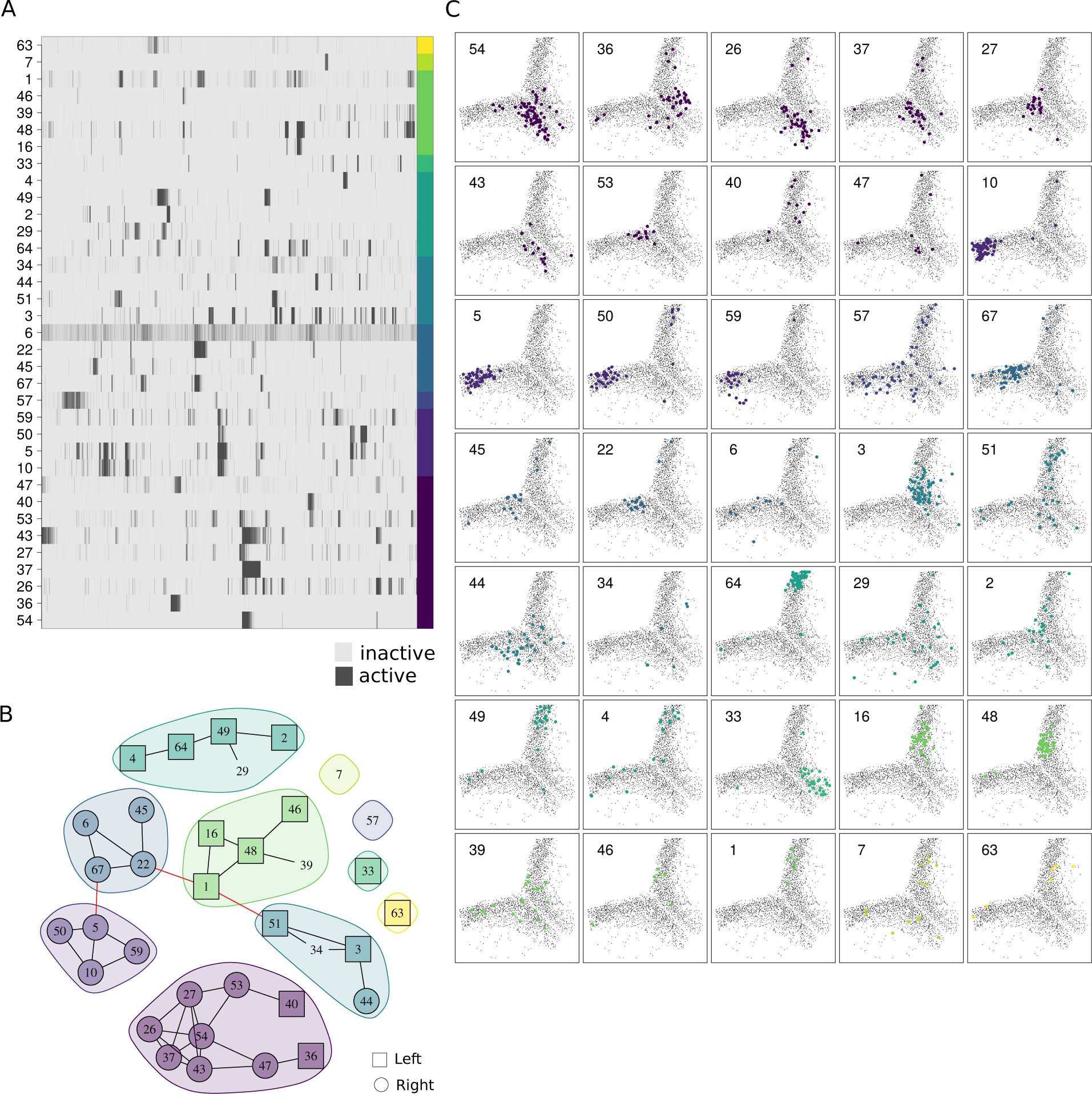
Distribution of tectal ensembles. (A) Neuronal ensembles from one of the images (different from the one presented in the main text) and the corresponding graph (B) representing the (Pearson’s) time correlation between ensembles.

**Figure S4:**
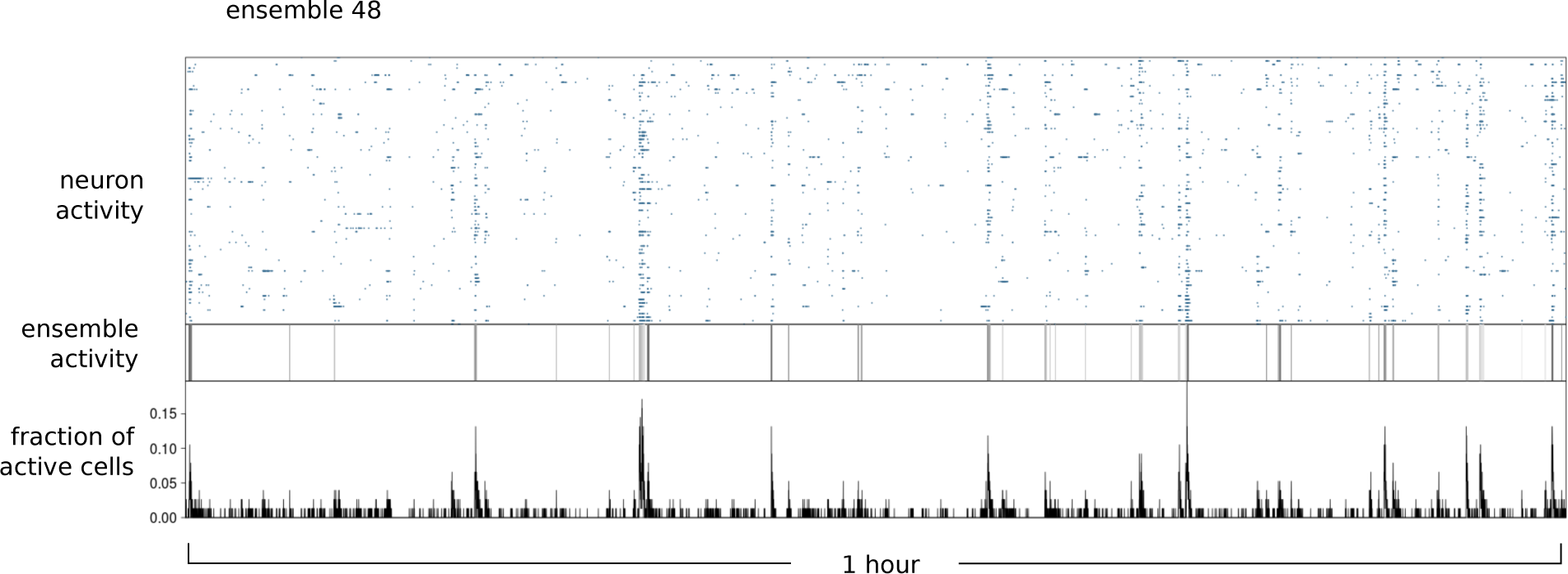
Ensemble vs cell activity. Comparison between ensemble activity and neuronal activity for the tectal ensemble 48 in Fig. 6.

**Figure S5:**
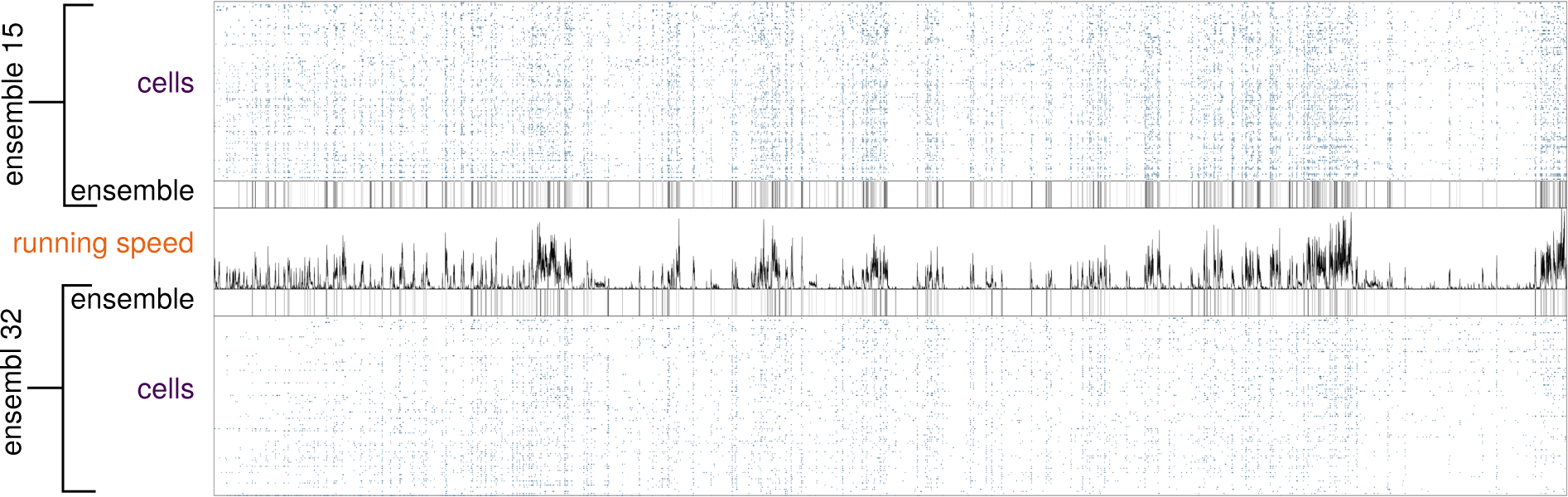
Differences between highly correlated ensembles. Comparison between ensembles 15 and 32 from Fig. 7.

